# Effects of Frontoparietal Theta tACS on Verbal Working Memory: Behavioral and Neurophysiological Analysis

**DOI:** 10.1101/2021.07.11.451931

**Authors:** Zhenhong Hu, Immanuel B.H. Samuel, Sreenivasan Meyyappan, Ke Bo Chandni Rana, Mingzhou Ding

**Affiliations:** J. Crayton Pruitt Family Department of Biomedical Engineering, University of Florida Gainesville, Florida, 32611

**Keywords:** tACS, Frontoparietal network, Verbal working memory, Working memory capacity, Theta oscillations, State dependent, Individual difference

## Abstract

Left-lateralized frontoparietal theta oscillations are thought to play an important role in verbal working memory. We causally tested this idea by stimulating the frontoparietal theta network at individual theta frequencies (4 to 8 Hz) during verbal working memory and observing the subsequent behavioral and neurophysiological effects. Weak electric currents were delivered via two 4×1 HD electrode arrays centered at F3 and P3. Three stimulation configurations, including in-phase, anti-phase, or sham, were tested on three different days in a cross-over design. On each test day, the subject underwent three experimental sessions: pre-, during- and post-stimulation sessions. In all sessions, the subject performed a Sternberg verbal working memory task with three levels of memory load (load 2, 4 and 6), imposing three levels of cognitive demand. Analyzing behavioral, EEG, and pupillometry data from the post-stimulation sessions, we report three results. First, in-phase stimulation improved task performance only in subjects with higher working memory capacity (WMC) and under higher memory load (load 6). Second, in-phase stimulation enhanced frontoparietal theta synchrony during working memory retention only in subjects with higher WMC under higher memory loads (load 4 and load 6), and the enhanced frontoparietal theta synchronization is mainly driven by enhanced frontal→parietal theta Granger causality. Third, the pupil diameter was not different irrespective of whether the preceding stimulation was in-phase, anti-phase, or sham. These findings suggest that theta tACS effects on verbal working memory were load- and subject-dependent, rooted in tACS-induced changes in frontoparietal network interactions, and not driven by changes in arousal levels.

## Introduction

Working memory (WM) is a cognitive system where information is held online temporally in service of behavioral goals. Neuroimaging and lesion studies have provided ample evidence that WM is supported by regions in frontal and parietal cortices (Chein and Schneider, 2005; Jonides et al., 2008; Owen et al., 2005). In particular, the central executive component of WM is linked to the frontal cortex (D’Esposito et al., 1995; Kane and Engle, 2002), whereas the storage component is associated with the parietal cortex (Champod and Petrides, 2010; Olson and Berryhill, 2009; Postle et al., 2006). In verbal working memory (VWM), in which the information remembered is language-related, there is further evidence suggesting a left-hemisphere dominance in these cognitive operations (D’Esposito et al., 1998; Smith et al., 1996).

Frontal and parietal regions interact during WM. These interactions are thought to be mediated by theta (4 – 8 Hz) oscillations (Buzsáki, 1996; Rutishauser et al., 2010; Sarnthein et al., 1998). Increased frontoparietal long-range theta synchrony accompanies increased demands on central executive functions in WM (Sauseng et al., 2005). Mathematically, synchrony measures trial-by-trial consistency of phase difference, but it does not specify the phase difference per se. According to the neuronal communication via neuronal coherence (NCNC) hypothesis (Fries, 2005), the theta phase difference between frontal and parietal sites is functionally significant, with the phase difference close to 0 degree (in-phase) or close to 180 degree (anti-phase) associated with facilitation or hindrance of neuronal communications. Consistent with this idea, Polanía et al. (2012) applied 6 Hz tACS over left prefrontal and parietal regions with either 0 degree relative phase (in-phase condition) or 180 degree relative phase (anti-phase condition) in a delayed letter discrimination task, and found that exogenously induced frontoparietal theta synchronization (in-phase stimulation) or desynchronization (anti-phase stimulation) significantly improves or degrades visual memory-matching performance as compared to sham stimulation. More recently, using a change detection task with images of real-world objects, Reinhart and Nguyen, (2019) applied in-phase tACS to prefrontal and temporal regions simultaneously in older adults, and found that it can bias frontotemporal functional connectivity, facilitate the neural integration, and enhance working-memory performance. To what extent in-phase theta tACS applied to frontoparietal cortex can facilitate verbal WM and modulate the underlying oscillatory network? We addressed this question in this study.

Frontoparietal theta is state-dependent and exhibits significant individual variability. Theta is higher in more cognitively demanding conditions (Jensen and Tesche, 2002; Payne and Kounios, 2009) and theta modulations by experimental conditions are stronger in individuals with stronger executive functions (Zakrzewska and Brzezicka, 2014). Testing the state-dependent effects of tACS, Violante et al. (2017) applied theta tACS to exogenously modulate oscillatory activity in right frontoparietal network in a visual WM task, and demonstrated that externally induced synchronization improved performance only when cognitive demands were high. Testing the subject-dependent effects of tACS, Tseng et al., (2018) applied 6 Hz tACS to modulate theta oscillation between the left and right parietal cortex with either in-phase, anti-phase, or sham sinusoidal current stimulation in a visual WM task, and found that in-phase theta tACS improved visual WM performance only in low-performers, with high-performers suffering a marginally significant visual WM impairment. We sought to test the state- and subject-dependent effects of frontoparietal theta tACS on verbal WM in this study.

tACS induces synaptic changes via spike-timing dependent plasticity (Vossen et al., 2015; Zaehle et al., 2010). Neural synchrony plays an important role in the support and promotion of synaptic plasticity (Bergmann and Born, 2018; Fell and Axmacher, 2011; Gregoriou et al., 2009; Wang, 2010). While theta tACS has shown promising behavioral effects, the neurophysiological underpinnings of these behavioral outcomes are less clear. Moreover, when comparing active vs sham stimulation, it is important to ascertain that the observed effects are related to modulations in the targeted neural circuit rather than to some nonspecific effects such as changes in arousal levels. We sought to shed light on these issues in this study.

We applied in-phase, anti-phase, and sham stimulation protocols to modulate frontoparietal theta in subjects performing a verbal WM task with three levels of WM load, and measured behavior, pupillometry, and EEG data in post-stimulation sessions. Individual differences in executive functioning were assessed using working memory capacity (WMC) in a separate experiment. We predicted that under higher WM loads and in subjects with better executive functioning, in-phase theta tACS would enhance frontoparietal theta synchrony and improve behavioral performance.

## Materials and Methods

### Participants

The experimental protocol was approved by the University of Florida Institutional Review Board. Twenty-five healthy volunteers (14 females, 23.28 ± 3.22 years of age) gave written informed consent and participated in the study. All subjects reported having no implanted electronic devices, no metal implants in the head, and no history of psychiatric or neurological disorders; they were also not current users of psychoactive medication, were not pregnant, and had normal or corrected-to-normal vision. Five participants did not complete all the study sessions and were therefore excluded from further analysis. The data from the remaining 20 subjects (11 females, 23.55 ± 3.35 years of age) were analyzed and reported here.

### Experimental Procedure

#### Study Design

As illustrated in Figure 1, the experiment employed a single-blind, cross-over, and sham-controlled design. It consisted of four study sessions with 1 week between consecutive sessions. During the first study session (baseline), participants took the OSPAN test online, which yielded working memory capacity (WMC). A 12-minute resting state EEG, comprising 6-minutes eyes-open rest and 6-minutes eyes-closed rest, were then recorded. Subsequently, the subjects performed the WM task for 30 minutes while their EEG and pupil data were recorded. For each of the following three study sessions, participants started with a pre-stimulation EEG and pupil data recording (30 minutes) in which they performed the verbal WM task (pre-stimulation session). Then, they performed the WM tasks for 30 minutes (during-stimulation session) while receiving in-phase, anti-phase, or sham theta tACS stimulation with the order of stimulation schemes randomized and counterbalanced across subjects. The stimulation session was followed by another 30 minutes of the WM task while EEG and pupil data were collected (post-stimulation session).

**Figure 1.**
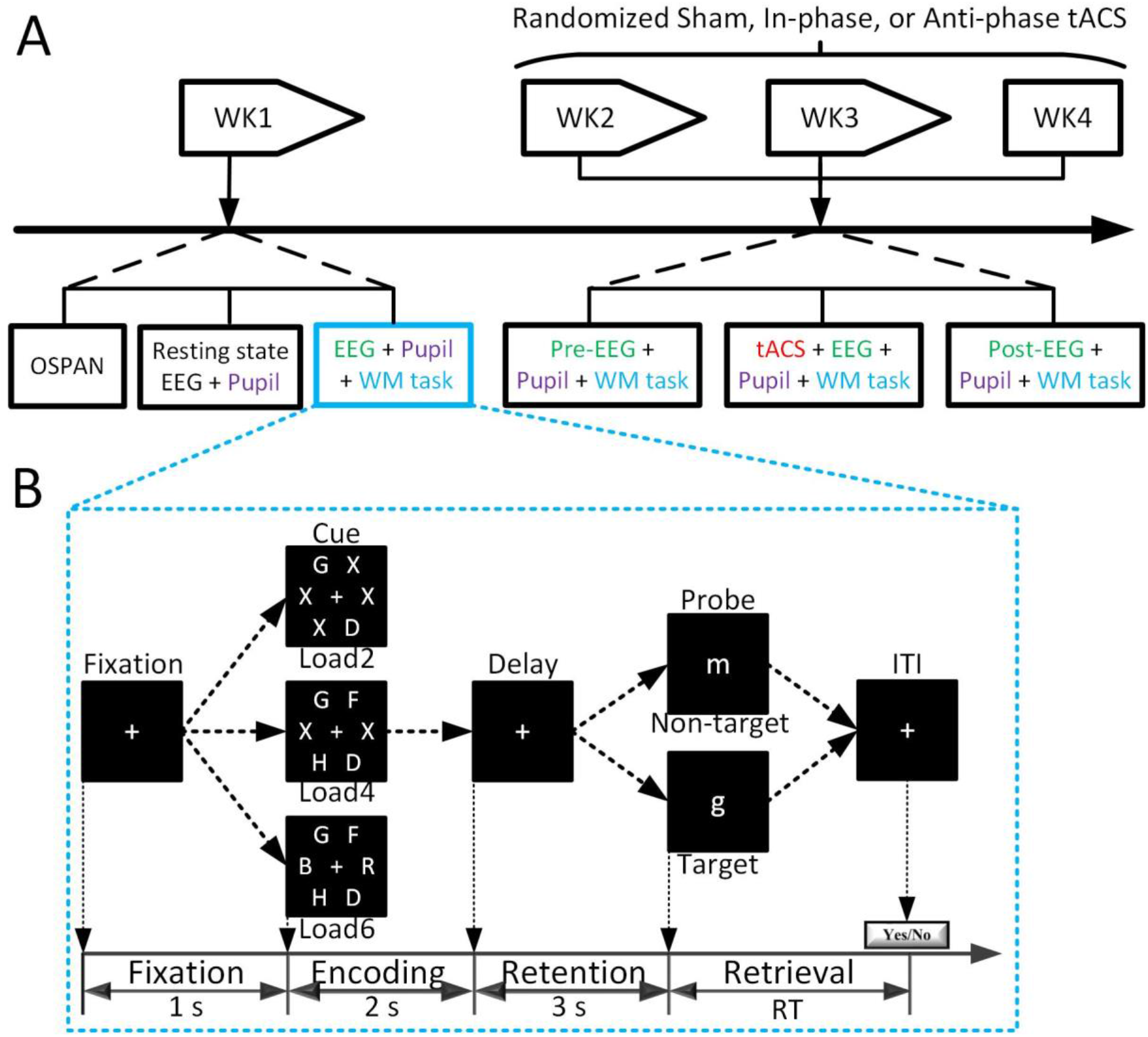
Overall experimental design and verbal working memory task. (A) The randomized, single-blind, cross-over, and sham-controlled design. (B) Timeline of the verbal working memory task with three levels of memory load (load 2, load 4, and load 6).

#### Verbal WM Task

In each of the four study sessions, participants performed a Sternberg WM task (Figure 1B). In this task, each trial started with a fixation cross presented at the center of the screen for 1 s, followed by a 2 s presentation of the memory set, which contained two, four, or six uppercase consonant letters placed with equal probability in any of six positions arranged in a circle centered on the fixation cross. When the memory set consisted of less than 6 letters, filler symbols (X) were added as a placeholder to make the sensory input for the three memory-load conditions comparable. The memory set varied randomly from trial to trial. The offset of the memory set was followed by a 3-second delay (retention), after which a lower-case letter, the probe, was shown at the center of the screen for 1 second. Subjects responded via a button press to indicate whether this character was part of the previously presented memory set. On half of the trials, the probe letter was part of the memory set, and on the other half, it was not. The filler symbol x was never used as a probe. Subjects were encouraged to respond as quickly and accurately as possible. The entire task consisted of three blocks with 72 trials in each block. The three memory loads were equally likely to occur. Breaks were given between blocks. Participants received a practice session prior to the experiment to familiarize with the task.

#### Administration of tACS

The tACS was administered using a Soterix Medical 1×1 HD-tES stimulator and two 4×1 HD-tES splitters. A schematic illustration of the electrode configuration was shown in Figure 2A. Five sintered mini Ag/AgCl electrodes attached to plastic holders filled with conductive gel were embedded in the Biosemi EEG cap to form each of the two 4×1 stimulation arrays. The center electrode of each array was placed at F3 and P3 respectively. The goal of the stimulation was to modulate synchronized neural oscillations in the theta band between the left frontal and left parietal cortex during verbal WM. Figure 2B shows the simulated electric field distribution (Soterix Medical HDExplore Software) associated with the in-phase HD-tACS protocols. Left frontal and left parietal cortex are maximally stimulated.

**Figure 2.**
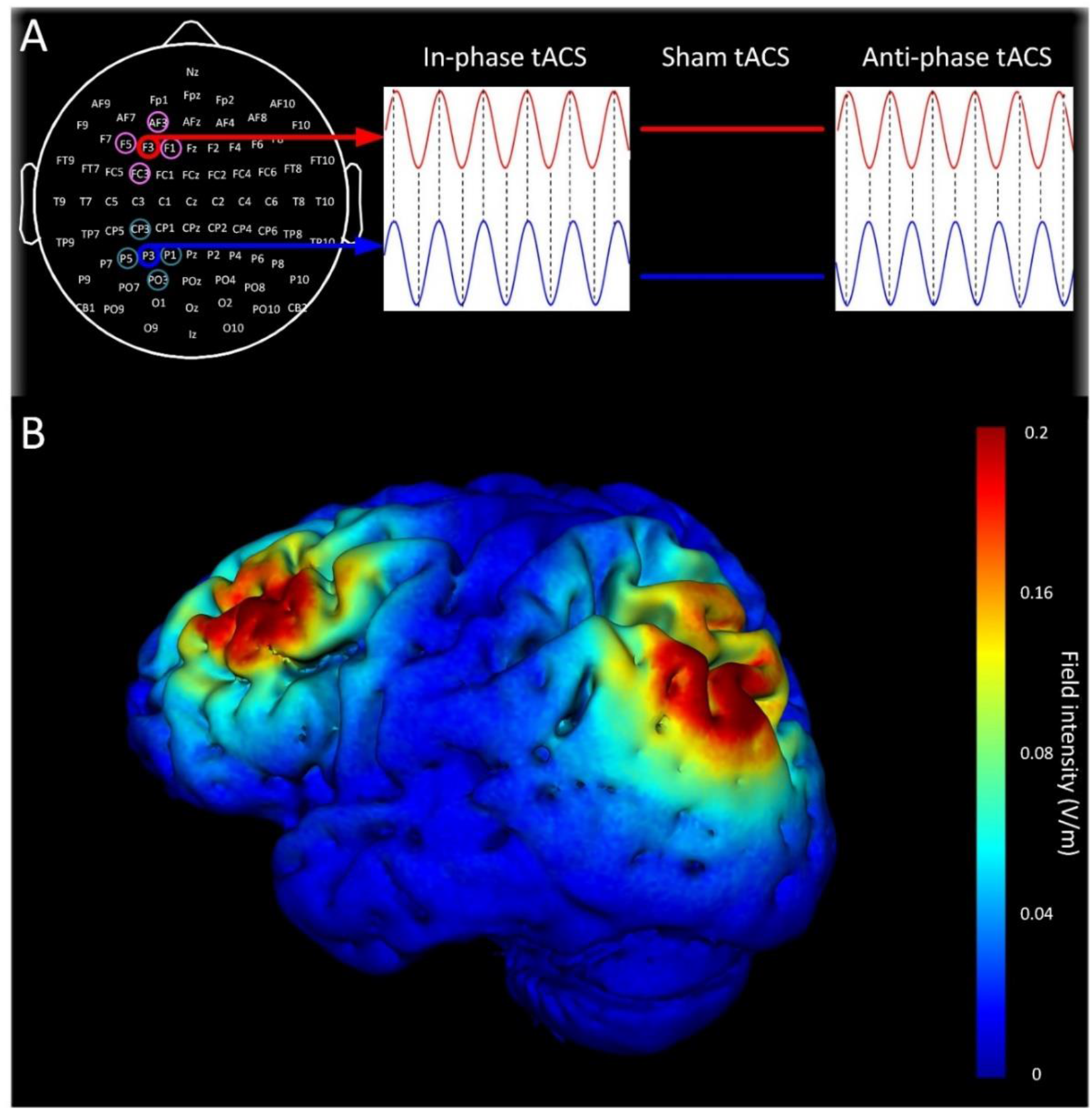
tACS arrays, stimulation schemes and simulated electric field distribution. (A) Position of the two 4×1 HD-tACS arrays and the three stimulation protocols: in-phase (0° phase difference), anti-phase (180° phase difference), and sham. Within each array, the center electrode and the four surrounding electrodes have opposite polarity, forming a closed circuit. The center-surround, source-sink arrangement of the five electrodes enables better focality of electrical stimulation (high definition). (B) Current flow shown on 3D reconstruction of the cortical surface demonstrates maximal electrical field intensity over the left frontal and parietal cortex.

Sinusoidal alternating current of 1 mA in magnitude was administered at each individual participant’s frontal peak theta frequency (PTF) for 30 min. The PTF, defined with a 0.5 Hz frequency resolution in the theta range (4-8 Hz), was determined from the EEG data recorded during Week 1 (WK1). Individual PTFs ranged from 4.50 - 6.50 Hz (M = 5.45, S.D. = 0.40 Hz) in the sample. All participants were familiarized with tACS-induced skin sensations with random noise stimulation of 30 s in duration. During tACS, the current ramped up to 1 mA over a time period of 30 s. In the in-phase condition, stimulation was delivered with 0° relative phase difference between two arrays, whereas in the antiphase condition, stimulation was delivered with a 180° relative phase difference. The sham stimulation condition followed the same procedure as the active condition, but stimulation only lasted 30 seconds, ramping up and down at the beginning and end of the 30-minute period, simulating the tingling sensation that subjects typically experience and then quickly habituate to during active stimulation sessions (Reinhart et al., 2017). For the pre-stimulation and post-stimulation sessions, the stimulating electrodes were replaced by EEG recording electrodes.

### Data Acquisition and Preprocessing

#### Data Acquisition

Experiments were performed in an electro-magnetically shielded room. Throughout the entire experiment, the subject’s pupil diameter was measured at a sampling rate of 1 kHz with an EyeLink 1000 infrared eye-tracker (SR Research, Mississaugu, ON, Canada). The subject’s EEG was recorded using a 128-channel Biosemi ActiveTwo system (Biosemi, Amsterdam, The Netherlands) at a sampling rate of 1 KHz.

#### Data Preprocessing

For continuous EEG data, the preprocessing was performed using EEGLAB (http://sccn.ucsd.edu/eeglab/index.html) and custom Matlab scripts (The Mathworks, Natick, MA, USA). The continuous EEG data were band-passed between 0.1 to 30 Hz, down-sampled to 256 Hz, and re-referenced against the average reference. For continuous pupil diameter data, blinks were detected using software provided by the manufacturer SR Research, and linear interpolation was carried out in Matlab. EEG and pupil data were epoched identically, from −1 s to 7 s with 0 s denoting the onset of the memory set (also referred to as cue). Trials with either excessive noise in the pupillary data or EEG were manually identified and removed. Trials with incorrect responses were also excluded from further analysis. For the remaining EEG trials, independent components analysis (ICA) (Delorme and Makeig, 2004) was applied to remove artifacts due to eye movements and blinks. For each memory load, the data from the middle 1 s of the retention period (3000-4000 ms) was selected as the time period of interest. Here, the first 1 s of the retention period, 2000-3000 ms, was excluded to avoid the negative impact of cue-offset-evoked activities on the spectral analysis of ongoing neural oscillations, and the last 1 s of the retention period, 4000-5000 ms, was excluded to avoid the negative impact of the anticipation of probe processing on neural activity (Wang and Ding, 2011). To minimize the negative effects of volume conduction and common reference on connectivity analysis (, the artifact-corrected scalp voltage data were converted to reference-free current source density (CSD) by calculating 2D surface Laplacian algorithm (Kayser and Tenke, 2006). All subsequent analyses were performed on the CSD data.

### Data Analysis

#### Working Memory Capacity

The participants’ individual WMC was assessed via the operation span task (OSPAN) (Unsworth et al., 2005) during Week 1. In each trial of this task, the subject was shown a series of letters to remember, and the number of letters to be remembered varied from 3 to 7 depending on the trial. A simple mathematical problem was inserted between letters, and at the end of the trial, the subject was asked to recall the letters from memory. There was a total of 15 trials. The OSPAN score, taken as a measure of WMC, was the sum of all correctly recalled letters across the 15 trials. The maximum OSPAN score is 75.

#### Theta Power Estimation

Fast Fourier transforms (FFT) were applied to the data in the time period of interest to estimate the power spectra. Normalization by power in the precue baseline period was done on a subject-by-subject basis (1 to 30 Hz) (Jensen et al., 2002). This normalization procedure removed the influence of amplitude variability from subject to subject and allowed more straightforward averaging across participants. Theta power were the averaged power from 4 to 8 Hz from the normalized power spectrum.

#### Phase Synchrony Estimation

We used phase locking value (PLV) (Lachaux et al., 1999) in theta frequency between two signals as a measure of neural synchrony. Specifically, the PLV at time *t* is defined as:

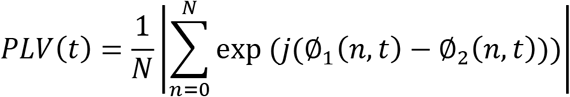

where *n* is the trial index, *N* = total number of trials, and the instantaneous phase value Ø_1_(*n,t*) and Ø_2_(*n,t*) in the theta band are extracted from the two signals using Hilbert transform. The PLV value measurers the inter-trial variability of the phase difference. PLV is close to 1 when the two signals are strongly coupled and close to zero if they are uncoupled. This procedure was repeated for all the pairwise channel combinations between the 5 frontal and 5 parietal recording channels.

#### Granger Causality (GC)

Neural synchrony measured by PLV was further decomposed into directional components using nonparametric GC (Dhamala et al., 2008; Ding et al., 2006). The nonparametric approach for estimating pairwise GC consists of the following steps: (i) using the multitaper method (Mitra and Pesaran, 1999) to construct spectral density matrix *S(f)* from Fourier transforms of two signals, (ii) factorizing spectral density matrix: *S* = ΨΨ* via Wilson’s algorithm (Wilson, 1972, 1978) where Ψ is the minimum-phase spectral factor, (iii) deriving noise covariance matrix ∑ and transfer function *H* from Ψ according to equations 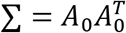 and 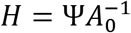, (iv) using *S,H*, and ∑ in Geweke’s formula (Geweke, 1982) to compute the causality from *y* to *x* at each frequency *f* according to:

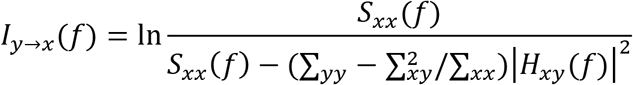

Reversing *y* and *x* in the above formula we can compute the causality from *x* to *y* at each frequency *f*.

## Results

### Working Memory Capacity

The average OSPAN across the subjects was 51.35 ± 14.75. Based on a median-split, the subjects were divided into a low WMC group and a high WMC group, with the low WMC group consisting of 6 women and 4 men (22.00 ± 2.62 years of age) and the high WMC group consisting of 5 women and 5 men (25.10 ± 3.38 years of age).

### TACS Effects on Task Performance

Over the entire sample of N=20 subjects, there were no significant differences in accuracy and response times for any of the three memory load conditions following in-phase, anti-phase, and sham stimulation (all *p* > 0.1). After splitting subjects into low and high WMC groups, we found that under the high memory load (load 6) condition, the response time post in-phase stimulation was significantly faster than that post sham stimulation (*t*_9_ = −2.47, *p* = 0.035) in subjects with high WMC (Figure 3 left). In-phase tACS yielded no behavioral benefits for the low WMC group (Figure 3 right). No significant effects were observed following the anti-phase tACS.

**Figure 3.**
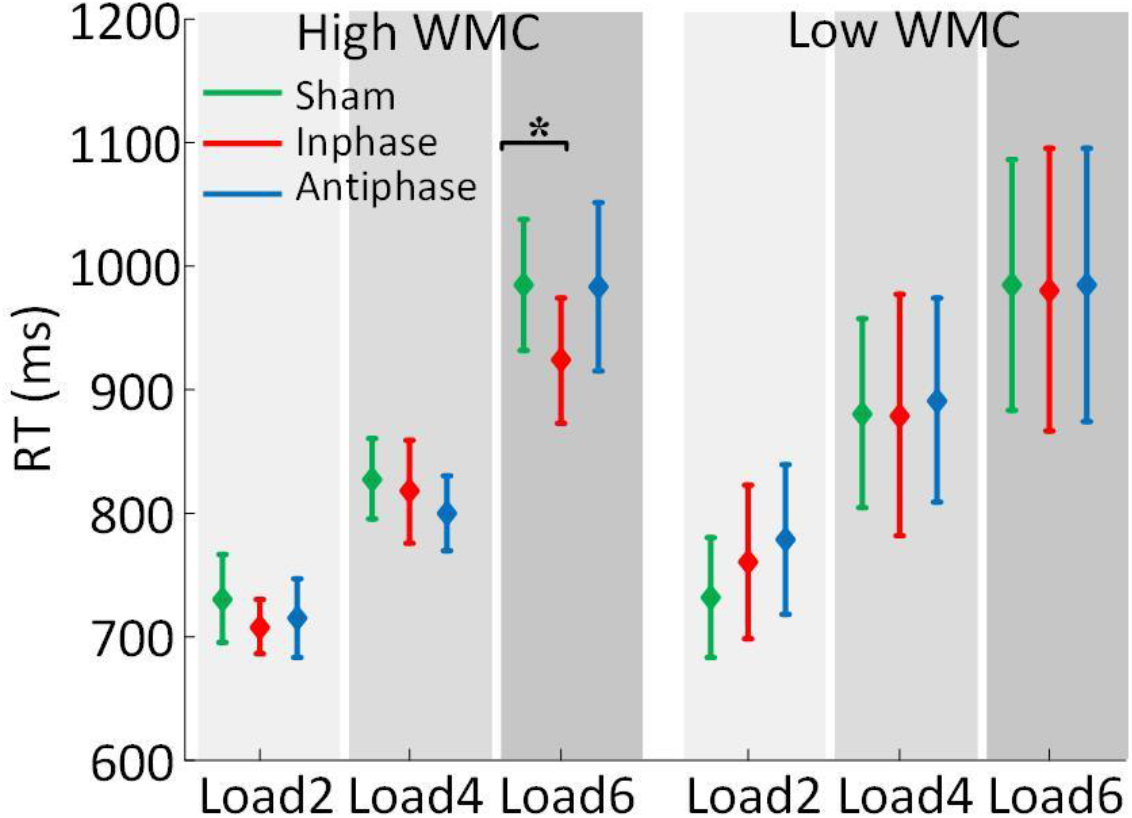
TACS effects on task performance. Mean RT under all WM load conditions for the three stimulation protocols in low (n = 10) and high (n = 10) WMC subjects. *p < 0.05. Green: post sham, red: post in-phase, blue: post anti-phase. *p<0.05.

### TACS Effects on Frontoparietal Theta Synchrony and Granger Causality

The interareal phase synchrony between frontal and parietal regions was assessed via PLV. There were no significant differences in theta PLV between left frontal and left parietal ROIs for any of the three memory load conditions across the three post stimulation sessions over the entire sample of N=20 subjects (all *p* > 0.1). However, after splitting subjects into low and high WMC groups, as shown in Figure 4A (top panel), we found that in subjects with high WMC, in-phase stimulation enhanced left frontoparietal theta synchronization relative to both sham and antiphase stimulation during working memory retention under memory load 4 and 6 (load 4: in-phase > sham, *t*_9_ = 3.12, *p* = 0.012; in-phase > antiphase, *t*_9_ = 2.60, *p* = 0.029. load 6: in-phase > sham, *t*_9_ = 2.44, *p* = 0.037; in-phase > antiphase, *t*_9_ = 2.71, *p* = 0.024). There was no evidence of enhanced theta synchronization (a) in low WMC individuals (Figure 4A, bottom panel) and (b) in the right frontoparietal network for either of the two WMC groups (Figure 4B).

**Figure 4.**
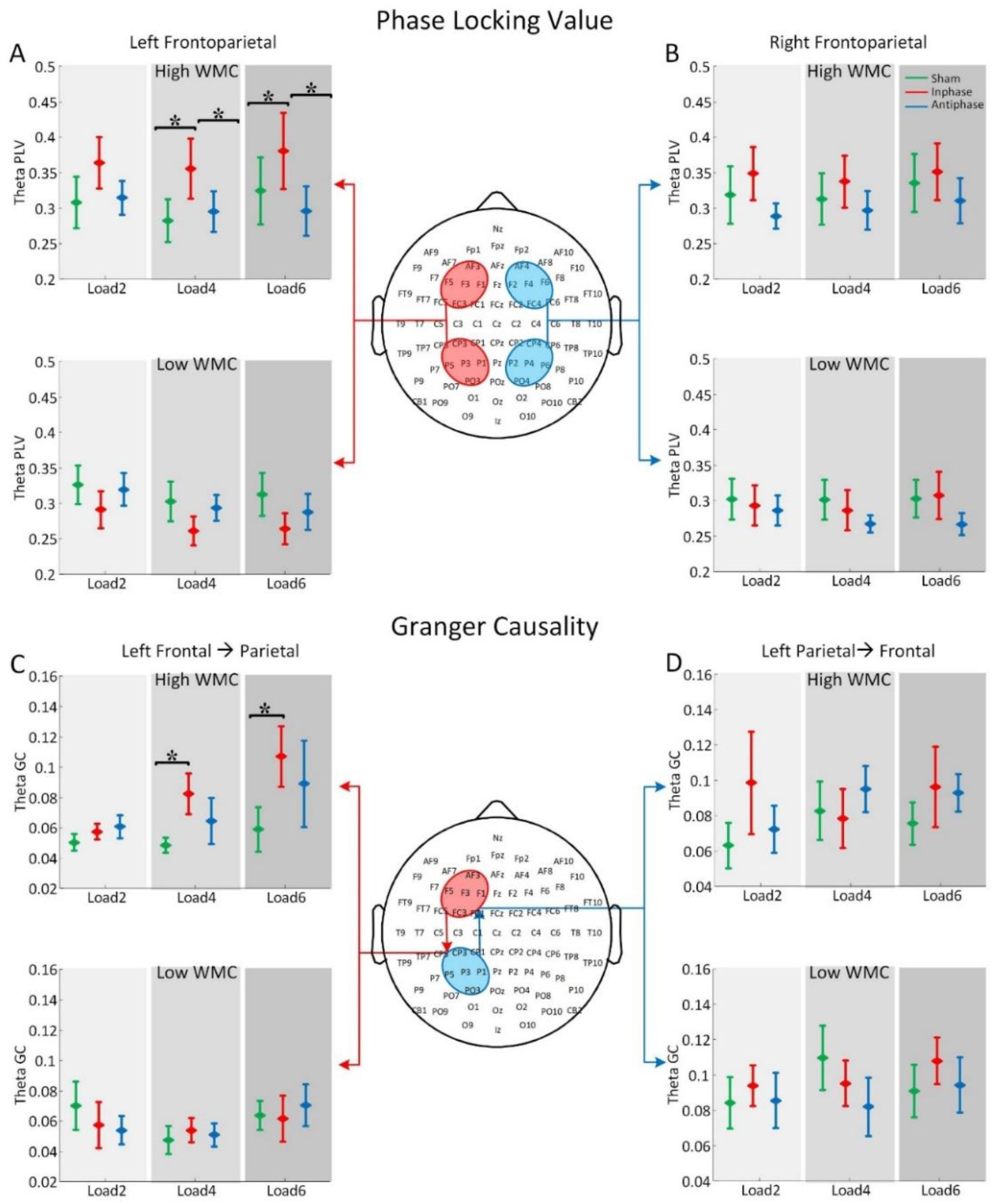
TACS effects on frontoparietal theta synchrony and Granger causality (GC). (A) Left frontoparietal theta phase locking value (PLV) for high and low WMC subjects. (B) Right frontoparietal theta PLV for high and low WMC subjects. (C) Left frontal → parietal theta band GC in high and low WMC subjects. (D) Left parietal → frontal theta band GC in high and low WMC subjects. Green: post sham, red: post in-phase, blue: post antiphase. *p<0.05.

Next, applying Granger causality (GC), the frontoparietal theta synchronization in the left hemisphere was decomposed into its directional components, frontal →parietal and parietal→frontal. As shown in Figure 4C, in-phase stimulation enhanced left frontal→parietal theta GC as compared to sham stimulation during WM retention under memory loads 4 and 6 in subjects with high WMC (load 4: in-phase > sham, *t*_9_ = 2.33, *p* = 0.045; load 6: in-phase > sham, *t*_9_ = 4.01, *p* = 0.0031). In contrast, there was no evidence of increased frontal→parietal theta GC in low WMC individuals (Figure 4C, bottom panel), and there was no difference in parietal→frontal theta GC in ether groups (Figure 4D). Thus, the increased frontoparietal synchrony in the theta band following in-phase stimulation under higher memory load conditions in high WMC individuals is mainly driven by increased frontal→parietal theta drive, whereas parietal →frontal GC is not modulated by theta tACS.

### TACS Effects on Pupil Diameter

Pupil diameter was examined to assess whether different stimulation schemes differentially affected arousal levels. As shown in Figure 5, there were no significant differences in pre-cue pupil diameter among the three post-stimulation sessions (all *p* > 0.1), indicating that the arousal level remained the same following sham, in-phase, and anti-phase stimulations. This result suggested that the observed effects on task performance, left frontoparietal theta synchronization, and left frontal → parietal theta GC were not due to changes in arousal levels.

**Figure 5.**
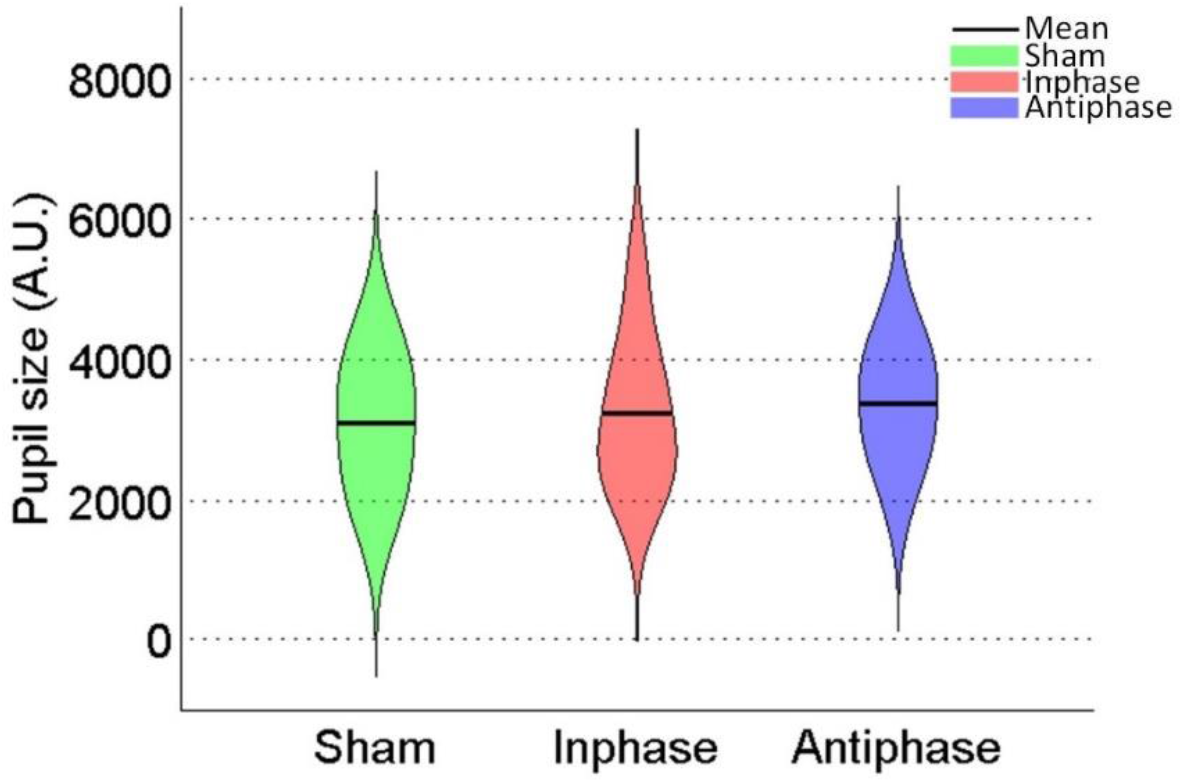
TACS effects on pupil size in the three post-stimulation sessions.

### Summary of TACS Effects

We summarized the findings reported in far in Table 1 where a check mark indicated that for that observable there was a significant difference post in-phase stimulation compared to other stimulation schemes. From the table it is clear that the effects were only observed under more demanding cognitive conditions in subjects with stronger executive functions.

**Table 1.** Summary of in-phase tACS effects

|  | Low WMC |  |  | High WMC |  |  |
| --- | --- | --- | --- | --- | --- | --- |
|  | Load2 | Load4 | Load6 | Load2 | Load4 | Load6 |
| Accuracy |  |  |  |  |  |  |
| RT |  |  |  |  |  | ✓ |
| Frontoparietal theta synchrony (l) |  |  |  |  | ✓ | ✓ |
| Frontoparietal theta synchrony (r) |  |  |  |  |  |  |
| Frontal → parietal theta GC (l) |  |  |  |  | ✓ | ✓ |
| Parietal → frontal theta GC (l) |  |  |  |  |  |  |
| Pupil diameter |  |  |  |  |  |  |

l: left and r: right.

## Discussion

We applied in-phase, anti-phase, and sham theta tACS to the left frontal (F3) and parietal (P3) sites via two 4×1 high-definition stimulation arrays during verbal working memory. In subjects with higher WMC and under more cognitively demanding conditions, in-phase theta tACS (1) improved WM task performance and (2) enhanced left frontoparietal theta synchrony and frontal→parietal theta Granger causality. The pupil diameter, an established marker of arousal, was found to be not different between stimulation schemes.

Both frontal and parietal cortices are key neural substrate supporting WM (Cohen et al., 1997; Pessoa et al., 2002; Todd and Marois, 2004). It is further suggested that theta oscillations provide the means for these regions to communicate during WM processes (Buzsáki, 1996; Rutishauser et al., 2010; Sarnthein et al., 1998). According to the neuronal communication via neuronal coherence (NCNC) hypothesis (Fries, 2005), in-phase oscillation between distant sites of an oscillatory network facilitates neuronal communication, whereas anti-phase oscillation hinders it. The advent of tACS, especially high definition tACS, has provided a technique to causally test these ideas. Polanía et al. (2012) applied 6 Hz tACS over left prefrontal and parietal regions with either 0 degree relative phase (in-phase condition) or 180 degree relative phase (anti-phase condition) in a delayed letter discrimination task, and found that exogenously induced frontoparietal theta synchronization (in-phase stimulation) or desynchronization (anti-phase stimulation) significantly improves or degrades visual memory-matching performance when compared to sham stimulation. Reinhart and Nguyen, (2019) applied in-phase tACS to target prefrontal and temporal regions simultaneously in older adults and found that it can bias frontotemporal-network connectivity and enhance working-memory performance. Our results extended these findings into verbal WM, and demonstrated that the effects were nuanced, exhibiting both state- and subject-dependence.

The magnitude of theta oscillations and synchrony is a function of the cognitive state and exhibits significant subject-to-subject variability. More cognitively demanding tasks are associated with stronger theta oscillations and synchrony. Individuals with stronger executive functioning show stronger theta modulation by cognitive conditions. It is thus expected that the effects of theta tACS vary across individuals and cognitive conditions. Alagapan et al., (2016) demonstrated in computational models that the modulation of cortical oscillations by brain stimulation depended on the brain state. Violante et al. (2017) applied theta tACS to exogenously modulate oscillatory activity in the right frontoparietal network in a visual n-back WM task, and demonstrated that externally induced synchronization improved performance when cognitive demands were high (2-back) and had no effect in the less demanding 1-back condition. Tseng et al. (2018) applied 6 Hz tACS to modulate theta oscillation between the left and right parietal cortex with either in-phase, anti-phase, or sham stimulation in a visual WM task and found in-phase theta tACS improved visual WM performance, but only in low-performers. Our results, showing that in-phase frontoparietal stimulation enhanced verbal working memory performance only in subjects with stronger executive functioning and under higher cognitively demand conditions, extend the idea that theta tACS is state- and subject-dependent, and demonstrate that it is more likely to be effective when there is a better developed theta oscillatory network and a stronger need for the involvement of theta activity.

Past tACS studies have mainly focused on the behavioral outcomes of stimulation. The neurophysiological underpinnings of these behavioral outcomes are less clear. We sought to shed light on this issue by recording both EEG and pupillometry data in the post-stimulation session where the stimulation electrodes were replaced by recording electrodes so that neural activity under the stimulation sites could be fully assessed. In our data, in-phase stimulation enhanced frontoparietal theta synchrony during working memory retention, and such enhancement was only observed in the left hemisphere (the stimulated hemisphere) but not in the right hemisphere, suggesting that the stimulation indeed led to enhancement in neuronal communication in the stimulated frontoparietal network. Similar to behavioral improvement, enhanced neuronal communications were only observed in individuals with stronger executive functioning and under higher cognitive demands, indicating that tACS did not alter the intrinsic properties of theta oscillation but the brain’s ability to better utilize these properties to accomplish cognitive tasks. Given that theta is an endogenous brain oscillation, having a better developed theta oscillatory system, as is the case in subjects with stronger executive functions, is the basis for benefiting from the effects of tACS stimulation. Decomposing neural synchrony into their directional components, Granger causality further revealed that the increased theta synchrony comes from increased frontal → parietal influence, whereas parietal →frontal influence remained unchanged, suggesting that improved executive control underlies improved verbal WM performance.

Can any of the changes we see following different tACS stimulation schemes be explained by nonspecific effects such as changes in the brain’s overall arousal levels? We used pupillometry data to address this question. Extensive research has shown that pupil size is a good indicator of the level of arousal and cognitive efforts (Aston-Jones and Cohen, 2005; Bradley et al., 2008; Ebitz and Platt, 2015; Eldar et al., 2013; Nassar et al., 2012; Urai et al., 2017). Since visual stimuli and cognitive loads are known to cause event-related pupil changes, to assess non-task related arousal levels, we focused the measurement of pupil diameter during the time period prior to the presentation of memory cues. Our results showed that the pupil size remained roughly the same in the post-stimulation session whether the session was preceded by in-phase, anti-phase, and sham stimulations. This can be taken as evidence to support the notion that the observed behavioral and neurophysiological effects following in-phase tACS were mainly due to changes in the frontoparietal network dynamics, not due to arousal level differences caused by different stimulation protocols.

It is worth noting that, although the NCNC hypothesis, when applied to network level stimulation predicts that: in-phase vs anti-phase stimulation should result in enhanced vs hindered neuronal communication and behavioral performance, empirically, not all studies have observed the predicted effects. Kleinert et al. (2017) applied 5 Hz tACS at fronto-parietal sites during a visuospatial match-to-sample task and reported that there were no significant differences between in-phase and anti-phase stimulation in both behavioral and EEG measurements. Miyaguchi et al., (2019) applied tACS at 70 Hz over the left M1 and the right cerebellar hemisphere in a visuomotor control task and found that the anti-phase stimulation decreased task error compared to the sham condition but did not differ from the in-phase stimulation. Violante et al. (2017) found a decrease in reaction time for in-phase tACS relative to sham and anti-phase tACS but did not observe any difference between anti-phase tACS and sham (Violante et al., 2017). In the present study, while we observed that in-phase stimulation enhanced theta synchrony and behavioral performance, there is no evidence of declining theta synchrony and task performance following anti-phase stimulation. The reasons for these discrepancies remain to be further elucidated in future studies.

Our study is not without limitations. First, EEG data during tACS stimulation were not analyzed because effectively separating neural activity from stimulation artifacts was difficult. It is thus not known whether different tACS protocols directly modified frontoparietal theta synchrony. This limitation is mitigated to some extent by recent studies showing that external tACS is capable of entraining and modulating endogenous brain oscillations in a frequency-specific manner (Feurra et al., 2011; Pogosyan et al., 2009; Polanía et al., 2012; Reinhart, 2017; Thut et al., 2011; Violante et al. 2017). Second, while the use of HD stimulations arrays improves stimulation focality compared to sponge-style electrodes, scalp-mounted devices still suffer from the lack of very precise spatial targeting ability. Third, the sample size is relatively small. One mitigating factor is that different from many previous studies where cross-sectional designs were utilized, we used a cross-over design, which is known to reduce variability, thereby permitting the observation of true effects in relatively small samples.

## Acknowledgments

This work was supported by the National Institute of Health grants MH112206.

## References

Alagapan, S., Schmidt, S.L., Lefebvre, J., Hadar, E., Shin, H.W., and Fröhlich, F. (2016). Modulation of Cortical Oscillations by Low-Frequency Direct Cortical Stimulation Is State-Dependent. PLOS Biology 14, e1002424.

Aston-Jones, G., and Cohen, J.D. (2005). An integrative theory of locus coeruleus-norepinephrine function: adaptive gain and optimal performance. Annu. Rev. Neurosci. 28, 403–450.

Bergmann, T.O., and Born, J. (2018). Phase-Amplitude Coupling: A General Mechanism for Memory Processing and Synaptic Plasticity? Neuron 97, 10–13.

Bradley, M.M., Miccoli, L., Escrig, M.A., and Lang, P.J. (2008). The pupil as a measure of emotional arousal and autonomic activation. Psychophysiology 45, 602–607.

Buzsáki, G. (1996). The Hippocampo-Neocortical Dialogue. Cereb Cortex 6, 81–92.

Champod, A.S., and Petrides, M. (2010). Dissociation within the frontoparietal network in verbal working memory: a parametric functional magnetic resonance imaging study. The Journal of Neuroscience 30, 3849–3856.

Chein, J.M., and Schneider, W. (2005). Neuroimaging studies of practice-related change: fMRI and meta-analytic evidence of a domain-general control network for learning. Cognitive Brain Research 25, 607–623.

Cohen, J.D., Perlstein, W.M., Braver, T.S., Nystrom, L.E., Noll, D.C., Jonides, J., and Smith, E.E. (1997). Temporal dynamics of brain activation during a working memory task. Nature 386, 604.

D’Esposito, M., Detre, J.A., Alsop, D.C., Shin, R.K., Atlas, S., and Grossman, M. (1995). The neural basis of the central executive system of working memory. Nature 378, 279.

Dhamala, M., Rangarajan, G., and Ding, M. (2008). Analyzing information flow in brain networks with nonparametric Granger causality. NeuroImage 41, 354–362.

Ding, M., Chen, Y., and Bressler, S.L. (2006). Granger Causality: Basic Theory and Application to Neuroscience. ArXiv:Q-Bio/0608035.

Ebitz, R.B., and Platt, M.L. (2015). Neuronal activity in primate dorsal anterior cingulate cortex signals task conflict and predicts adjustments in pupil-linked arousal. Neuron 85, 628–640.

Eldar, E., Cohen, J.D., and Niv, Y. (2013). The effects of neural gain on attention and learning. Nat. Neurosci. 16, 1146–1153.

d’Esposito, M., Aguirre, G.K., Zarahn, E., Ballard, D., Shin, R.K., and Lease, J. (1998). Functional MRI studies of spatial and nonspatial working memory. Cognitive Brain Research 7, 1–13.

Fell, J., and Axmacher, N. (2011). The role of phase synchronization in memory processes. Nature Reviews Neuroscience 12, 105–118.

Feurra, M., Bianco, G., Santarnecchi, E., Testa, M.D., Rossi, A., and Rossi, S. (2011). Frequency-Dependent Tuning of the Human Motor System Induced by Transcranial Oscillatory Potentials. J. Neurosci. 31, 12165–12170.

Fries, P. (2005). A mechanism for cognitive dynamics: neuronal communication through neuronal coherence. Trends in Cognitive Sciences 9, 474–480.

Geweke, J. (1982). Measurement of Linear Dependence and Feedback Between Multiple Time Series. Journal of the American Statistical Association 77, 304–313.

Gregoriou, G.G., Gotts, S.J., Zhou, H., and Desimone, R. (2009). High-Frequency, Long-Range Coupling Between Prefrontal and Visual Cortex During Attention. Science 324, 1207–1210.

Jensen, O., and Tesche, C.D. (2002). Frontal theta activity in humans increases with memory load in a working memory task. European Journal of Neuroscience 15, 1395–1399.

Jonides, J., Lewis, R.L., Nee, D.E., Lustig, C.A., Berman, M.G., and Moore, K.S. (2008). The mind and brain of short-term memory. Annu Rev Psychol 59, 193–224.

Kane, M.J., and Engle, R.W. (2002). The role of prefrontal cortex in working-memory capacity, executive attention, and general fluid intelligence: An individual-differences perspective. Psychonomic Bulletin & Review 9, 637–671.

Kayser, J., and Tenke, C.E. (2006). Principal components analysis of Laplacian waveforms as a generic method for identifying ERP generator patterns: I. Evaluation with auditory oddball tasks. Clinical Neurophysiology 117, 348–368.

Kleinert, M.-L., Szymanski, C., and Müller, V. (2017). Frequency-Unspecific Effects of θ-tACS Related to a Visuospatial Working Memory Task. Front Hum Neurosci 11.

Lachaux, J.-P., Rodriguez, E., Martinerie, J., and Varela, F.J. (1999). Measuring phase synchrony in brain signals. Human Brain Mapping 8, 194–208.

Mitra, P.P., and Pesaran, B. (1999). Analysis of dynamic brain imaging data. Biophys J 76, 691–708.

Miyaguchi, S., Otsuru, N., Kojima, S., Yokota, H., Saito, K., Inukai, Y., and Onishi, H. (2019). Gamma tACS over M1 and cerebellar hemisphere improves motor performance in a phase-specific manner. Neuroscience Letters 694, 64–68.

Nassar, M.R., Rumsey, K.M., Wilson, R.C., Parikh, K., Heasly, B., and Gold, J.I. (2012). Rational regulation of learning dynamics by pupil–linked arousal systems. Nat Neurosci 15, 1040–1046.

Olson, I.R., and Berryhill, M. (2009). Some surprising findings on the involvement of the parietal lobe in human memory. Neurobiology of Learning and Memory 91, 155–165.

Owen, A.M., McMillan, K.M., Laird, A.R., and Bullmore, E. (2005). N-back working memory paradigm: A meta-analysis of normative functional neuroimaging studies. Human Brain Mapping 25, 46–59.

Payne, L., and Kounios, J. (2009). Coherent oscillatory networks supporting short-term memory retention. Brain Research 1247, 126–132.

Pessoa, L., Gutierrez, E., Bandettini, P.A., and Ungerleider, L.G. (2002). Neural Correlates of Visual Working Memory: fMRI Amplitude Predicts Task Performance. Neuron 35, 975–987.

Pogosyan, A., Gaynor, L.D., Eusebio, A., and Brown, P. (2009). Boosting Cortical Activity at Beta-Band Frequencies Slows Movement in Humans. Curr Biol 19, 1637–1641.

Polanía, R., Nitsche, M.A., Korman, C., Batsikadze, G., and Paulus, W. (2012). The Importance of Timing in Segregated Theta Phase-Coupling for Cognitive Performance. Current Biology 22, 1314–1318.

Postle, B.R., Ferrarelli, F., Hamidi, M., Feredoes, E., Massimini, M., Peterson, M., Alexander, A., and Tononi, G. (2006). Repetitive Transcranial Magnetic Stimulation Dissociates Working Memory Manipulation from Retention Functions in the Prefrontal, but not Posterior Parietal, Cortex. Journal of Cognitive Neuroscience 18, 1712–1722.

Reinhart, R.M. (2017). Disruption and rescue of interareal theta phase coupling and adaptive behavior. Proceedings of the National Academy of Sciences 201710257.

Reinhart, R.M.G., and Nguyen, J.A. (2019). Working memory revived in older adults by synchronizing rhythmic brain circuits. Nat Neurosci 22, 820–827.

Reinhart, R.M.G., Cosman, J.D., Fukuda, K., and Woodman, G.F. (2017). Using transcranial direct-current stimulation (tDCS) to understand cognitive processing. Atten Percept Psychophys 79, 3–23.

Rutishauser, U., Ross, I.B., Mamelak, A.N., and Schuman, E.M. (2010). Human memory strength is predicted by theta-frequency phase-locking of single neurons. Nature 464, 903–907.

Sarnthein, J., Petsche, H., Rappelsberger, P., Shaw, G.L., and von Stein, A. (1998). Synchronization between prefrontal and posterior association cortex during human working memory. Proc Natl Acad Sci U S A 95, 7092–7096.

Sauseng, P., Klimesch, W., Schabus, M., and Doppelmayr, M. (2005). Fronto-parietal EEG coherence in theta and upper alpha reflect central executive functions of working memory. International Journal of Psychophysiology 57, 97–103.

Smith, E.E., Jonides, J., and Koeppe, R.A. (1996). Dissociating verbal and spatial working memory using PET. Cerebral Cortex 6, 11–20.

Thut, G., Schyns, P.G., and Gross, J. (2011). Entrainment of Perceptually Relevant Brain Oscillations by Non-Invasive Rhythmic Stimulation of the Human Brain. Front Psychol 2.

Todd, J.J., and Marois, R. (2004). Capacity limit of visual short-term memory in human posterior parietal cortex. Nature 428, 751.

Tseng, P., Iu, K.-C., and Juan, C.-H. (2018). The critical role of phase difference in theta oscillation between bilateral parietal cortices for visuospatial working memory. Scientific Reports 8, 349.

Unsworth, N., Heitz, R.P., Schrock, J.C., and Engle, R.W. (2005). An automated version of the operation span task. Behavior Research Methods 498–505.

Urai, A.E., Braun, A., and Donner, T.H. (2017). Pupil-linked arousal is driven by decision uncertainty and alters serial choice bias. Nat Commun 8, 14637.

Violante, I.R., Li, L.M., Carmichael, D.W., Lorenz, R., Leech, R., Hampshire, A., Rothwell, J.C., and Sharp, D.J. (2017). Externally induced frontoparietal synchronization modulates network dynamics and enhances working memory performance. ELife 6, e22001.

Violante, I.R., Li, L.M., Carmichael, D.W., Lorenz, R., Leech, R., Hampshire, A., Rothwell, J.C., and Sharp, D.J. Externally induced frontoparietal synchronization modulates network dynamics and enhances working memory performance. ELife 6.

Vossen, A., Gross, J., and Thut, G. (2015). Alpha Power Increase After Transcranial Alternating Current Stimulation at Alpha Frequency (α-tACS) Reflects Plastic Changes Rather Than Entrainment. Brain Stimulation 8, 499–508.

Wang, X.-J. (2010). Neurophysiological and computational principles of cortical rhythms in cognition. Physiol Rev 90, 1195–1268.

Wang, X., and Ding, M. (2011). Relation between P300 and event-related theta-band synchronization: A single-trial analysis. Clinical Neurophysiology 122, 916–924.

Wilson, G. (1972). The Factorization of Matricial Spectral Densities. SIAM J. Appl. Math. 23, 420–426.

Wilson, G.T. (1978). A convergence theorem for spectral factorization. Journal of Multivariate Analysis 8, 222–232.

Zaehle, T., Rach, S., and Herrmann, C.S. (2010). Transcranial Alternating Current Stimulation Enhances Individual Alpha Activity in Human EEG. PLOS ONE 5, e13766.

Zakrzewska, M.Z., and Brzezicka, A. (2014). Working memory capacity as a moderator of load-related frontal midline theta variability in Sternberg task. Frontiers in Human Neuroscience 8.

